# Faf2 is required for neural differentiation in embryonic neural progenitor cells

**DOI:** 10.64898/2026.07.12.737973

**Authors:** Anneke Dixie Kakebeen, Leon Dunphy, Heather K Hazen, Lee A Niswander

**Affiliations:** Department of Molecular, Cellular, and Developmental Biology, University of Colorado Boulder

**Keywords:** Neural differentiation, neural progenitor cells, FAF2, ER stress

## Abstract

Neural progenitor cell differentiation is a complex process requiring the proper integration of instructive and permissive factors. Instructive cues including signaling molecules and transcription factor networks have been well studied in this context, but permissive factors such as cell homeostasis have not. Cell homeostasis is critical to support the health and stability of a cell and enable the cell to act on instructive differentiation cues. Our study investigates a homeostasis protein, FAF2, and its function in neural progenitor cells. FAF2 is an adaptor protein involved in endoplasmic reticulum (ER) associated degradation to remove misfolded proteins and restore ER homeostasis. Here we show that knocking out *Faf2* in neural progenitor cells results in increased ER stress signature at the protein and transcription level, indicating a conserved functional role in neural progenitor cells. Induced neural differentiation of FAF2 deletion cells shows a failure of neurite development but RNA-seq indicates genes that support neural differentiation are induced. Reducing ER stress in FAF2 knockout cells with a small molecule inhibitor can rescue neural differentiation, providing evidence that excess ER stress contributes to the inhibited differentiation. Taken together, these results reveal that FAF2 is a critical protein in neural progenitor cells for the maintenance of ER homeostasis and execution of neural differentiation.

**Highlights:**

- FAF2 is required to regulate ER homeostasis in neural progenitor cells
- FAF2 knockout blocks differentiation of neural progenitor cells to neurons at the cell morphological level, but does not inhibit the mounting of transcriptional programs associated with neural differentiation.
- Excess ER stress due to FAF2 knockout contributes to blocked neural differentiation.

## Introduction

Neural progenitor cells are an essential component of the developing embryonic central nervous system. These cells must divide to expand the progenitor pool while also gaining specificity along their developmental trajectory to ultimately form the great diversity of neuronal subtypes and differentiate into mature neurons. Cell fate decisions are driven by a myriad of cell intrinsic and extrinsic factors (Bonnefont and Vanderhaeghen, 2021; Wang et al., 2025a, 2025b). These factors can be instructive, directing cell fate down a specific trajectory, or permissive, allowing cells to respond to the instructive cues (Tatapudy et al., 2017). Imbalance of these factors and failure of neural progenitor cells to execute the correct cell fate decisions can be disruptive to the generation of the nervous system. Thus, it is critical to gain a wholistic model of the types of cues that govern neural progenitor cell fate decisions.

Morphogen signaling and downstream transcription factor networks are instructive cues that have been well studied in neural progenitor cell fate. Removing these factors inhibits the cell from mounting the necessary cascade of signals to commit the cell toward a specific fate. Permissive cues provide the correct cellular environment to respond to instructive cues and carry out the cell fate changes. Permissive cues include processes that support cell stability and survival such as cell homeostasis and metabolism. While there is some evidence that permissive cues are necessary to support cell fate decisions in neural progenitor cells (Tatapudy et al., 2017), the links between cell homeostasis and the ability to respond to and carry out instructive cues and the homeostatic proteins that are integral to promoting differentiation is still understudied.

Endoplasmic reticulum (ER) stress can be a source of cellular instability (Rutkowski and Kaufman, 2004; Spencer and Finnie, 2020). The ER is the primary site of protein folding and quality control. When there is a shift in the ER toward more misfolded proteins, the ER enters a stressed state and employs strategies to regain homeostasis such as initiating the unfolded protein response (UPR). UPR serves to reduce the burden of misfolded proteins by reducing translation and increasing machinery that can remove misfolded proteins and chaperone the folding of new proteins (Hetz et al., 2020; Rutkowski and Kaufman, 2004; Zhang and Kaufman, 2006). The UPR functions through the activation of three receptors (ATF6, PERK, IRE1) which undergo modifications when the UPR chaperone, BIP, no longer binds to these receptors (Bertolotti et al., 2000; Hetz et al., 2020). The activated receptors induce transcription factor localization to the nucleus to turn on transcriptional programs aimed at reducing the burden of misfolded proteins.

ER stress and increased unfolded protein response are hallmark phenotypes in neurodegenerative disease and some neurodevelopmental contexts (Godin et al., 2016; Kawada et al., 2014; Murao and Nishitoh, 2017; Saito and Imaizumi, 2018; Wu et al., 2024). An increase in protein aggregates is often present in neurodegenerative diseases and can overwhelm the ER, sending the UPR into full affect (Saito and Imaizumi, 2018). In neurodevelopment, several neural tube defect models show increased ER stress as part of the pathobiology (Li et al., 2013; Yan et al., 2021). Although it is known that ER stress can modulate nervous system development, there is no consensus model of how ER stress can influence neural stem and progenitor cell fate decisions. In some cases, inducing the UPR by increasing ER stress can increase cell differentiation, in others too much ER stress can inhibit differentiation, and in some cases differentiation can proceed but with defects in neuronal maturation (Godin et al., 2016; Kawada et al., 2014; Vásquez et al., 2022). Several ER-stress associated proteins such as Sel1L, Hrd1, and ATF4 have been studied in the context of neural stem cell fate decisions and shown to play critical roles (Frank et al., 2010; Kawada et al., 2014; Saito et al., 2023). Even so, there is still limited information of the role that homeostatic proteins play in cell fate decisions. The goal of our study is to test the context-specific effect of ER stress on embryonic neural progenitor cell fate through the investigation of a homeostasis protein.

Fas-associated factor 2 (FAF2, UBXD8, ETEA) is an ER membrane protein that functions as an adaptor to extract misfolded proteins from the ER to the cytoplasm for degradation (Mueller et al., 2008). FAF2 contains a ubiquitin associated (UBA) domain which identifies ubiquitinated proteins and a ubiquitin regulatory x (UBX) domain which recruits the ATPase, p97/VCP, to extract the ubiquitinated protein for proteasomal degradation (Christianson et al., 2011; Mueller et al., 2008; Zheng et al., 2022) (**Figure 1A)**. FAF2 also modulates lipid biology by targeting proteins involved in lipid synthesis and lipid droplet regulation (Ganji et al., 2023, p. 97; Olzmann et al., 2013; Suzuki et al., 2012) and FAF2 is important for regulation of peroxisome homeostasis (Kim et al., 2025). Although these functions have been studied across several different cell types (HeLa, HEK293, U2OS, hepatocytes, adult neurons (Huda et al., 2025; Van Alstyne et al., 2025; Zheng et al., 2022), the function of FAF2 has not been studied in the context of stem cells, and specifically neural stem and progenitor cells. Here we establish FAF2 as a homeostasis protein in embryonic neural progenitor cells that is required for neuronal differentiation.

**Figure 1.**
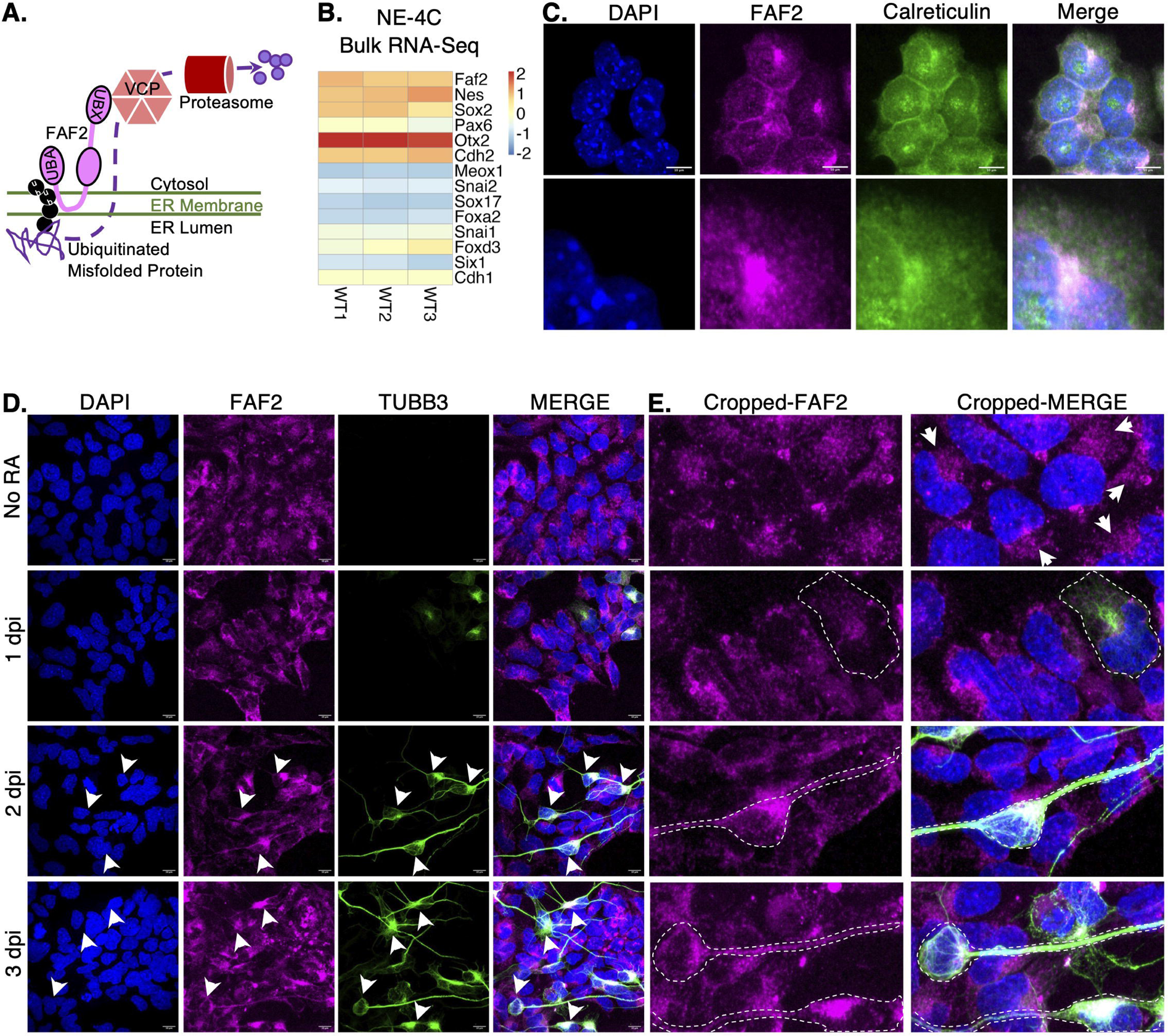
FAF2 is expressed in mouse embryonic neural progenitor cells. A) Schematic of FAF2 domain structure and role in ER-regulated degradation of misfolded proteins. UBA: Ubiquitin-associated domain; UBX:ubiquitin regulatory x domain; Ub: ubiquitin. B) Heatmap showing relative expression of genes that mark the three germ layers derived from bulk RNA-Seq of individual replicates of wildtype NE-4C cells. C) Immunofluorescence of wildtype NE-4C cells against FAF2 (magenta) and Calreticulin (ER marker in green). Cells were counterstained with DAPI to visualize the nucleus. D) Immunofluorescence time-course of retinoic acid (RA) induced neural differentiation in wildtype NE-4C cells against FAF2 (magenta) and TUBB3 (neuronal marker in green). Arrows show differentiated neurons. Dpi = days post-induction. E) Zoomed images from (D).

## Materials and Methods

### NE-4C cell line

NE-4C cells were originally derived from E9 mouse embryonic brain (Schlett and Madarász, 1997). Our stocks were purchased from ATCC (CRL-2925). Cells were grown at 37°C and 5% CO_2_ conditions in MEM (Gibco 41090101) supplemented with 10% FBS, 1xMEM nonessential amino acids (Gibco 11140050), 1xGlutaMax (Gibco 35050061), and 1% penicillin-streptomycin (Gibco 15070063). The NE-4C cell line was routinely tested for mycoplasma contamination.

### Genome editing using CRISPR/Cas9

The CRISPR-Cas9 DNA editing method to generate the *Faf2* knockout cell line was derived from (Zocher and Kempermann, 2021). Two sgRNAs were used to create a 700bp deletion to remove exon 4 of *Faf2* and create an early stop codon We used sgRNA sequences (CCCACTGCGATGACCATGCA; CGATCATAACACCTTCCACT) as was done to generate the same deletion in a *Faf2* knockout mouse model by the Knockout Mouse Project (Faf2em1(IMPC)Mbp)(Wilson et al., 2026) (https://www.mousephenotype.org/). To create sgRNA/Cas9 plasmids we used the pSpCas9(BB)-2A-Puro (PX459) V2.0 plasmid from Addgene (62988). This plasmid contains the Cas9 coding sequence, a puromycin resistance cassette and cloning region for an sgRNA. Individual plasmids were cloned for each sgRNA sequence by inserting cloning-adapted oligos purchased from IDT into the *BbsI* cloning site as detailed in (Zocher and Kempermann, 2021) (sgRNA1: F-caccCCCACTGCGATGACCATGCA/R- aaacTGCATGGTCATCGCAGTGGG; sgRNA2:F- caccCGATCATAACACCTTCCACT/ R- aaacAGTGGAAGGTGTTATGATCG). Insertion of the sgRNA was verified by Sanger sequencing.

To edit cells, NE-4C cells were transfected with both sgRNA plasmids using Lipofectamine 3000 (ThermoFisher Scientific L3000015) one day after plating 200,000 cells in a 12-well plate. On day 2-post transfection, the media was replaced with selection media containing 3µg/mL puromycin for 2-4 days. After puromycin selection, remaining cells were re-plated to a 10cm plate and individual colonies were picked to a 24 well plate when colonies reached ∼100 cells. Colonies were expanded to maintain the stock and genomic DNA collected for PCR. Colonies showing only one band at 525bp representing the deletion product were then validated by Sanger sequencing.

### Knockdown cell line generation by shRNA

Two shRNA plasmids targeting *Faf2* were purchased from the University of Colorado Anschutz Functional Genomics Shared Resource (RRID: SCR_021987). Plasmids included the *Faf2* shRNA sequence and puromycin resistance gene for selection. NE-4C cells were transfected with two shRNA plasmids (TRCN0000191263: CGGTTTACCTATTACACGATA; TRCN0000191032: CCATGACTTCTTATTCTCCTT) by Lipofectamine 3000. On day 2-post transfection, the media was replaced with selection media containing 3µg/mL puromycin for one week. Remaining cells were kept as a poly-clonal line. To confirm random integration of the plasmids into the cells for stable expression of the shRNA, genomic DNA was collected and used for PCR. PCR products from a forward primer common to the U6 promoter of the shRNA plasmid and reverse primer specific to the target sequence at 250bp represent DNA specific to the introduced plasmids that was integrated into the cellular genome.

### Neuronal differentiation and small molecule treatments

NE-4C neuronal differentiation was induced by addition of 1µM all-trans retinoic acid (RA) (Sigma-Aldrich # 302-79-4) for 24 hours followed by incubation in growth media (Varga et al., 2008). To induce ER stress, NE-4C cells were treated with 0.01µM Thapsigargin (Tocris 1138) for 24 hours. Stressed cells were then either collected for immunofluorescence or thapsigargin media was replaced with differentiation media. To reduce ER stress, NE-4C cells were treated with 1mM 4-phenylbutyric acid (4-PBA, Cayman Chemical 11323) for 24 hours. For differentiation experiments, 4-PBA treated cells were then cultured in media containing 4-PBA and RA for 24 hours.

### Immunofluorescence, imaging, and analysis

For immunofluorescence experiments, cells were plated directly on glass coverslips in cell culture plates. Following experimentation, cells were fixed on coverslips in 4%PFA for 15 minutes at room temperature, washed in 1xPBS and then permeabilized in 0.1% Triton in 1xPBS (PBST) for 15 min at room temperature. Cells were blocked in PBST containing 1% normal goat serum from 30 minutes at room temperature before addition of primary antibodies diluted in blocking solution. Primary antibodies were incubated overnight at 4^0^C. Following primary antibody incubation, cells were washed with PBST, then secondary antibody (1:500) and DAPI diluted in PBST were added for 2 hours at room temperature. Cells were then washed in 1xPBS before being mounted on glass slides with Prolong Gold Antifade (Invitrogen P10144). Cells were imaged with a Yokogawa CellVoyager™ CV1000 spinning disk scanner housed by the University of Colorado Light Microscopy Core Facility (RRID:SCR_018993). Primary antibodies used: TUBB3 (BioLegend PRB-435P rabbit, 1:1000); FAF2 (ThermoFisher 66629-1-IG mouse, 1:300); pH3 (Cell Signaling Technology 9714 rabbit, 1:100); BIP (CST 3177S rabbit, 1:300); CALR (Abcam ab2907 1:100 rabbit).

### Thioflavin T staining

Cells were plated on glass coverslips and allowed to grow to ∼70-80% confluence. Media was then replaced with cell culture media containing 10µg/mL Hoechst to stain nuclei and 5µM Thioflavin T (Cayman Chemical 32553) to stain protein aggregates for 30 minutes in the 37°C, 5% CO_2_ incubator. Staining media was replaced with 1xPBS and coverslips were mounted on glass slides and immediately imaged on the Yokogawa CellVoyager™ CV1000 spinning disk scanner.

### Quantitative Image Analysis

For each imaging experiment, 5-10 fields of view were imaged per coverslip and experiments were repeated a minimum of three times using different cell passages. For FAF2 expression, thioflavin T and differentiation experiments, ImageJ was used to measure pixel intensity of the DAPI channel and experimental channel (FAF2, BIP, TUBB3, ThT) from thresholded images. Experimental channel intensities were normalized to DAPI area for a normalized intensity value. The normalized value was log2 transformed and means were calculated from fields of view per experiment. For proliferation experiments, the number of pH3 positive cells were counted and normalized to the number of nuclei identified by DAPI for five fields of view per biological replicate. The mean log2 values or pH3/cell for the fields of view per each biological replicate were used to statistically test for differences between conditions. For experiments with two conditions, a Student’s t-test was used to test for significance; for experiments with more than two conditions, a two-way Anova was run with a post-hoc Tukey test to derive pairwise p-values.

### Western Blots

Cells were collected by trypsinization, pelleted at 4000 RPM for 5 minutes, and washed in PBS before pelleting again. Cell pellets were resuspended in RIPA buffer and lysed on ice for 30 minutes. Lysate was then spun down at max speed for 20 minutes at 4^0^C and supernatant was taken. Cell lysates were quantified by Bradford assay. Lamelli buffer was added to protein lysates at a 1:1 ratio and boiled for 15 minutes.

15 µg total protein were loaded onto 7.5% polyacrylamide gels for anti-FAF2 blots and 4-20% gradient polyacrylamide gels for all other blots and run at 200V for 30 minutes. Protein was then transferred to PVDF membrane using the BioRad Turbotransfer system. Membranes were blocked in 5% milk in TBST for 30 minutes, then primary antibody was added to the blocking solution and incubated overnight at 4^0^C. Following washing, secondary HRP antibodies were added to blots in blocking solution at 1:2000 dilution and incubated at room temperature for 1 hour. Blots were washed and visualized on a BioRad Chemidoc with ECL substrate. Primary antibodies used: TUBB3 (BioLegend PRB-435P rabbit, 1:10000); FAF2 (ThermoFisher 16251-1-AP rabbit, 1:1000); pH3 (Cell Signaling Technology 9714 rabbit, 1:500); BIP (CST 3177S rabbit, 1:1000); p-eIF2a (CST 3398S rabbit, 1:1000); p-IRE1 (Fisher PA1-16927 rabbit, 1:000); ATF6 (Novus NBP175478SS rabbit, 1:1000); GAPDH (rabbit, 1:10000).

Western blots were quantified in FIJI using the gel analysis tool. Values for each protein per sample were normalized to the corresponding GAPDH band intensity. Log transformed normalized values were statistically compared and averages of minimum three biological replicates shown on plots. Experiments with two conditions were statistically tested by Student’s t-test; experiments with more than two conditions were statistically tested by an Anova with a post-hoc Tukey test for pairwise p-values.

### Bulk RNA-Seq preparation and processing

WT and knockout cells were collected from three independent passages and frozen as cell pellets. The “NO” condition (not induced) were collected without perturbations. The “RA” condition (retinoic acid induced) were treated for 24 hours with RA followed by basal media until collection at 3 days post induction. Total RNA was then extracted from cell pellets and sent for mRNA library preparation by The University of Colorado, Anschutz Shared Genomics Core (RRID: SCR_021984) using the Universal Plus™ mRNA-Seq library preparation kit with NuQuant. Libraries were sequenced at a depth of 40 million paired end reads on an Illumina NovaSeq.

Fastq files were processed and returned as a counts table using the established NF-Core RNA-Seq pipeline (https://nf-co.re/rnaseq/3.22.2/) in bash (Ewels et al., 2020). We used the default pipeline which uses STAR for alignment and Salmon for quantification. Reads were aligned to Mus_musculus.GRCm39.109. The output counts were normalized using DESeq2 in R using the vst command. We then used DESeq2 to perform differential expression analysis on desired comparisons (Love et al., 2014) (Full differential expression gene list in **Supplementary Table 1**). Raw Fastq files and metadata will be made available on SRA and GEO.

### GSEA and Gene Ontology analysis

Gene set enrichment analysis was performed using the R package fgsea (Korotkevich et al., 2021). Genes were ranked and ordered by the Wald statistic which is an output of differential expression analysis in DESeq2. The ranked list was inputted into the function *fgsea* and the GO:BP database for *Mus musculus* was queried. Only significantly enriched pathways are reported. Full gene set enrichment analysis results in **Supplementary Table 2**.

Gene ontology (GO) analysis for transcription factor enrichment was performed utilizing the R package gProfileR (Raudvere et al., 2019). Genes that were significantly increased or decreased in expression between WT and KO cells were treated separately as inputs for analysis. The GO terms from source “TF” were investigated for transcription factor binding motifs. The genes that called each transcription factor were then used for GO analysis; the output of this analysis was investigated for GO:Biological Processes. Full gene ontology analysis results in **Supplementary Tables 3/4**.

## Results

### FAF2 is present in embryonic mouse neural progenitor cells

While it is known that FAF2 is present in the ER of other cell types including hepatocytes, HeLa cells, HEK cells, and adult neurons, the role of FAF2 during embryonic neural development is unknown. To study FAF2 function in the context of embryonic neural stem cells, NE-4C cells were used as a model system. NE-4C cells are a commercially available cell line derived from embryonic day 9 (E9) mouse brain and immortalized by deletion of p53 (Schlett and Madarász, 1997). To provide a baseline for our studies and as a general resource, we performed RNA-Seq of WT NE-4C cells. Gene expression from WT NE-4C cells show that the neural progenitor cell molecular identity is preserved as revealed by higher expression of neural progenitor marker genes (*Nes, Sox2, Pax6, Otx2, Cdh2)* relative to genes that represent other cell types (*Meox1, Snai2, Sox17, Foxa2, Snai1, Foxd3, Six1, Cdh1)* (**Figure 1B**). *Faf2* is also expressed in neural progenitor cells and immunofluorescence confirmed FAF2 protein expression in NE-4C cells. As in other cell types, FAF2 has perinuclear localization and colocalizes with Calreticulin (CALR), which indicates its localization to the endoplasmic reticulum (**Figure 1C**).

### FAF2 protein is present during neural differentiation

Neural progenitor cells are specialized cells that must balance proliferation and differentiation to expand the progenitor pool while also producing new neurons. NE-4C cells are capable of differentiation by induction with All-Trans Retinoic Acid (RA) (Schlett and Madarász, 1997; Varga et al., 2008), thus we tested if FAF2 expression or localization differs between neural progenitor cells and neurons. For differentiation experiments, cells were treated with retinoic acid (RA) for 24 hours and then RA was washed out and chased with growth media. Cells were collected to visualize expression of FAF2 and TUBB3, an early neuronal marker, over the three days post induction (dpi). FAF2 expression was similar in cycling neural progenitor cells and differentiating neurons (**Figure 1D**) with FAF2 polarized to one side of the nucleus and found in the cell bodies of neural progenitors and neurons. FAF2 protein was also found in the neurites of differentiating neurons (**Figure 1D/E**).

### Loss of FAF2 results in ER stress signature

FAF2 is critical for modulating ER stress and promoting cell homeostasis in other cell types, however this role has not been investigated in neural stem cells. To test if FAF2 protein expression is responsive to ER stress, NE-4C cells were treated with Thapsigargin (TG) to induce ER stress. Thapsigargin is a small molecule that depletes the ER of calcium, leading to disrupted protein folding (Li et al., 1993). Cells were treated with thapsigargin for 18 hours and fixed for immunofluorescence with BIP, a chaperone of the unfolded protein response and used as an indicator of ER stress (Sicari et al., 2020). Thapsigargin treated cells had significantly more BIP pixel intensity than untreated cells (**Figure 2A/B**). FAF2 expression was also increased in Thapsigargin treated cells indicating FAF2 protein expression in neural progenitor cells is responsive to acute ER stress (**Figure 2A/B**).

**Figure 2.**
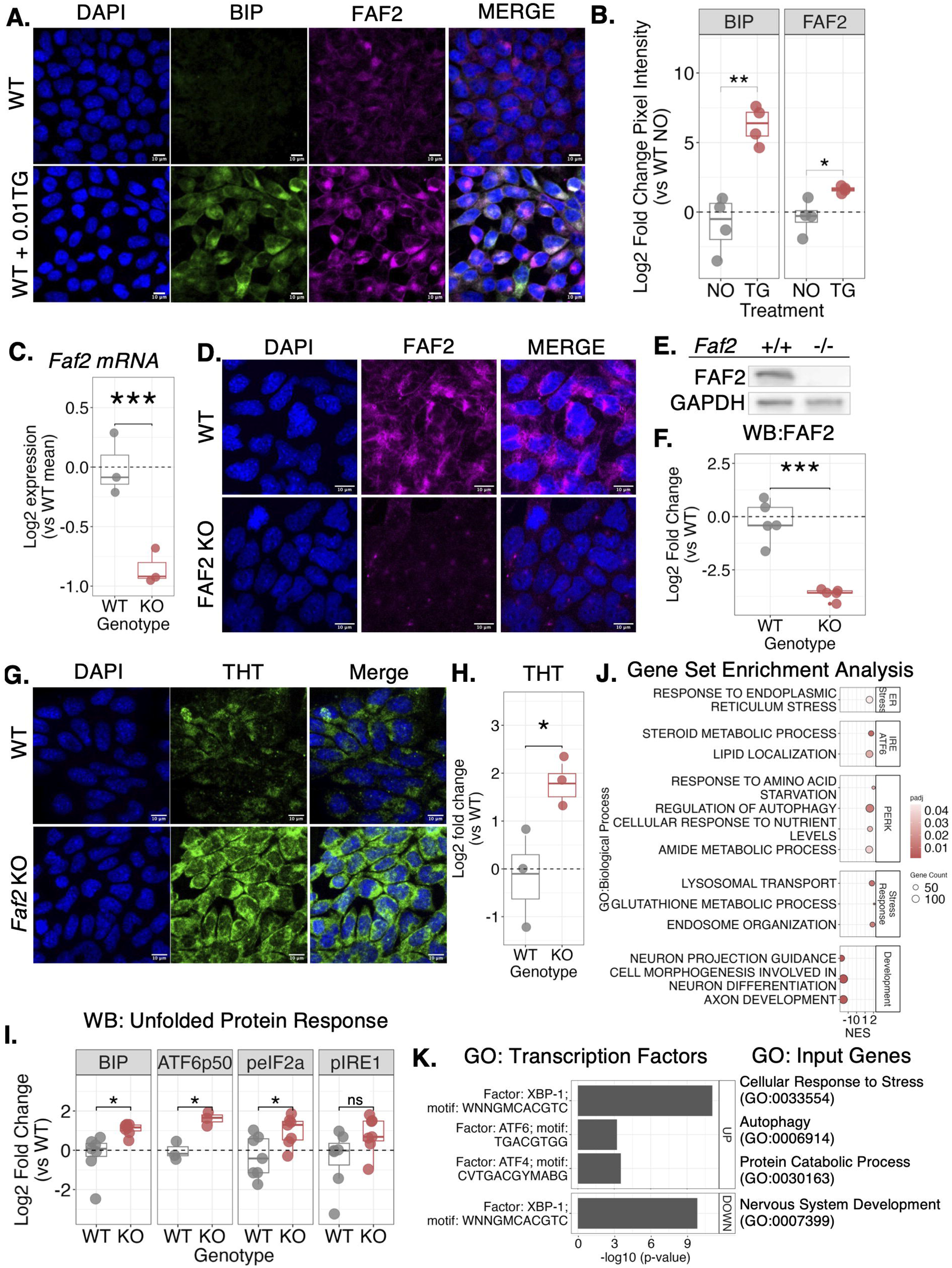
FAF2 knockout results in increased ER stress signature in neural progenitor cells. A) Immunofluorescence against BIP (ER stress marker in green) and FAF2 (magenta) in untreated and Thapsigargin (TG) treated wildtype cells. B) Quantification of pixel intensity of BIP and FAF2 as in (A). Each condition shows n=4 biological replicates. C) mRNA expression of *Faf2* in WT and *Faf2 KO* cells from bulk RNA-Seq data. D) Immunofluorescence images of FAF2 (magenta) in WT and *Faf2 KO* cells E) Western blot of WT and *Faf2 KO* cells to assess FAF2 levels compared to GAPDH. F) Quantification of western blots as in (E). n=5 biological replicates. G) Immunofluorescence images of thioflavin T (THT) staining (marker of misfolded proteins) of WT and *Faf2 KO* cells. Cells were counterstained with DAPI. H) Quantification of THT pixel intensity from (G). I) Quantification of western blots for unfolded protein response proteins. Representative western blot shown in Supplementary Figure 1D. J) Gene set enrichment analysis from bulk RNA-Seq of WT and *Faf2 KO* cells. Graph shows representative terms related to cellular stress. Full list of terms can be found in the supplement. K) Gene-ontology analysis results for transcription factor calls. “UP” represents transcription factors called from up-regulated genes and “Down” represents transcription factors called from down-regulated genes *p<0.05, **p<0.01, ***p<0.001

To test the hypothesis that FAF2 is required to reduce ER stress in neural progenitor cells, we generated a *Faf2* knockout line using CRISPR/Cas9 mediated exon 4 deletion. We identified successful deletion of exon 4 followed closely by a premature stop codon by PCR and Sanger sequencing (**Supplementary 1A/B**). *Faf2* mRNA expression was significantly reduced in KO cells (**Figure 2C).** The FAF2 protein has several functional domains necessary to perform its role as an adaptor to extract misfolded proteins from the ER for proteasomal degradation. Exon 4 overlaps with the hydrophobic domain necessary to anchor FAF2 to the ER membrane and the early stop codon is predicted to terminate translation in exon 5 which would exclude the downstream UAS and UBX domains (**Supplementary 1C**). Without the hydrophobic domain, FAF2 will not anchor to the ER and without the UBX domain, FAF2 will not be able to recruit p97/VCP which is required for protein dislocation (Olzmann et al., 2013; Suzuki et al., 2012). FAF2 protein expression was successfully knocked out from NE-4C deletion cells as shown by immunofluorescence and western blot (**Figure 2D-F**).

The *Faf2 KO* cell line was used to investigate whether the levels of various ER homeostasis markers changed in response to a loss of FAF2. FAF2 is important for the clearance of misfolded proteins and therefore it reduces the buildup of protein aggregates. To test the hypothesis the *Faf2* KO would lead to a buildup of protein aggregates, protein aggregates were visualized by Thioflavin T (ThT) staining in WT and KO cells. Thioflavin T is a small molecule dye that has fluorescent properties when bound to beta-sheets present in protein aggregates; thus its incorporation and imaging has been used as a way to monitor the buildup of misfolded proteins and ER stress (Beriault and Werstuck, 2013). Indeed, KO cells had significantly more ThT pixel intensity than untreated WT cells, suggesting that *Faf2* KO results in an increase in protein aggregation (**Figure 2G, H**).

Protein aggregation in the ER can trigger the unfolded protein response (UPR). The UPR is made up of three signaling branches that can be detected at the protein and transcription level (Sicari et al., 2020). Each branch (ATF6, PERK, IRE1,) signals through receptors and corresponding activated transcription factors to drive expression of genes responsible to resolve the conflict. To test if the KO cells showed a signature of the UPR, western blot was used to detect protein expression of markers of the UPR. BIP was significantly expressed in KO cells suggesting an overall engagement of the UPR (**Figure 2I, Supplementary 1D).** To evaluate the three branches, antibodies against ATF6, p-eIF2a/PERK, and p-IRE were used. ATF6p50 and p-eIF2a were significantly increased in KO cells suggesting engagement of the ATF6 and PERK branches of the UPR (**Figure 2I, Supplementary 1D**). p-IRE1 also showed increased expression, although this was not statistically significant (**Figure 2I, Supplementary 1D**).

To test if the KO cells showed a transcriptional signature of the UPR, RNA-Seq was used as an unbiased means to profile the transcriptome. Differentially expressed genes between WT and KO cells were scored using the Wald statistic in DESeq2. Using all scored genes regardless of significance, we ran a gene set enrichment analysis (GSEA) to identify overrepresented pathways. Pathways that were positively enriched in KO cells included terms representative of all three UPR branches (IRE1/ATF6: “Steroid Metabolic Process”, “Lipid Localization; PERK “Response to Amino Acid Starvation”, “Regulation of Autophagy”, “Cellular Response to Nutrient Levels”, “Amide Metabolic Process”) (**Figure 2J, full GSEA results in Supplementary Table 2**). Pathways that were negatively correlated with the KO cells represented processes having to do with neural development and differentiation (“Neuron Projection Guidance”, “Cell Morphogenesis Involved in Neuron Differentiation”, “Axon Development”) (**Figure 2J**).

Given the gene sets that were enriched in KO cells, we next asked if relevant UPR-associated transcription factor motifs were also enriched. Using gProfiler and the TRANSFAC database, we identified predicted transcription factor binding motifs enriched in KO cells. In this analysis, the input genes represent differentially expressed target genes of transcription factors. These genes have been annotated with the inclusion of transcription factor binding sites within 1kb of the transcription start site (Raudvere et al., 2019). Genes that showed increased expression in the KO cells revealed ATF4, ATF6, and XBP1 binding motifs (**Figure 2K**). ATF4 or ATF6 binding motifs were not represented in genes that were downregulated, however XBP1 motifs were called out for these genes (**Figure 2K).** gProfiler was used to identify gene ontology terms from the discrete set of genes which identified each transcription factor. The target genes of ATF4, ATF6, and XBP1 that were significantly increased in KO cells represented stress response terms consistent with the UPR pathways represented by each transcription factor **(Figure 2K, Supplementary Figure 2).** In contrast, the target genes that called XBP1 and were significantly decreased did not represent stress response genes, but instead called terms involved with developmental processes including the term “nervous system development” **(Figure 2K, Supplementary Figure 2, full Gene ontology results in Supplementary Table 4)**. Overall, our data provides evidence that knocking out FAF2 results in a signature of ER stress, thus demonstrating a requirement for FAF2 to maintain ER homeostasis in NE-4C neural progenitor context.

### FAF2 is required for neural differentiation

Since knocking out FAF2 showed a robust effect on ER stress modulation in neural progenitor cells, we next wanted to investigate if neural progenitor functions were sensitive to loss of FAF2. To test if cell proliferation rates were altered in KO cells, immunofluorescence and western blots against phospho-histone 3 (pH3) were used in WT and KO cells. Calculating the percent of pH3 positive cells per region of interest showed no significant difference between WT and KO cells, which was validated by western blot analysis of pH3 in whole protein lysates (**Figure 3A-C, H-I**).

**Figure 3.**
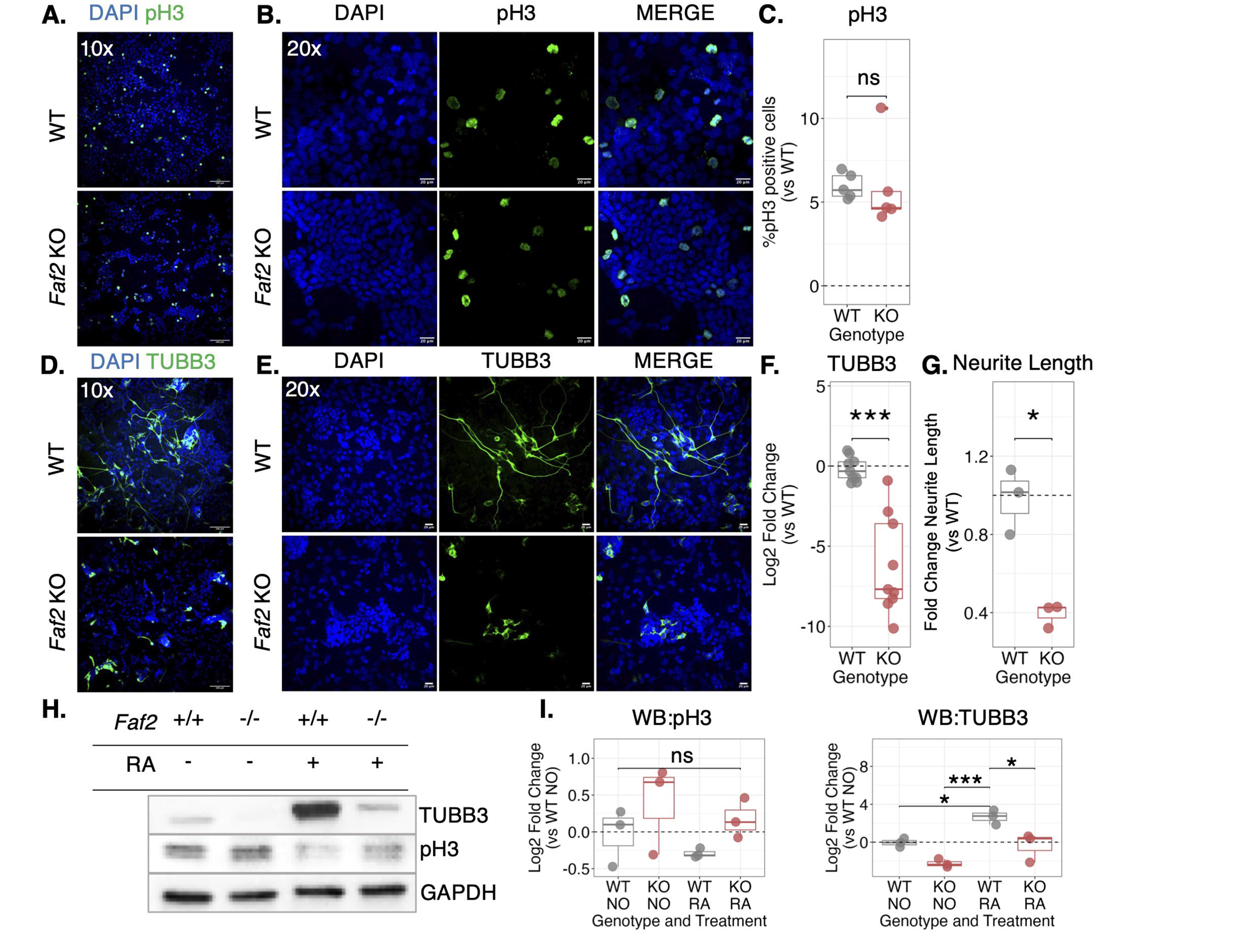
FAF2 is required for neural differentiation of NE-4C cells. A) Immunofluorescence images against pH3 in WT and *Faf2 KO* cells at 10x magnification, counterstained with DAPI. B) Representative field of view at 20x magnification of the same samples in (A). C) Quantification of the percent pH3 positive cells per field of view. 20x objective was used for collecting data for quantification. D) Immunofluorescence images against TUBB3 in WT and *Faf2 KO* cells at 10x magnification, counterstained with DAPI E) Representative field of view at 20x magnification of same samples in (D). F) Quantification of pixel intensity of TUBB3 relative to DAPI channel in (E). G) Quantification of neurite length in images from (E). 20x objective was used for collecting data for quantification in F and G. H) Western blot against TUBB3, pH3, and GAPDH in WT and *Faf2 KO* cells that were untreated or treated with RA to induce differentiation. I) Quantification of pH3 and TUBB3 bands from western blot as in (H). n = 3 biological replicates. Treatments: NO = untreated, RA = treated with all-trans retinoic acid. *p<0.05, **p<0.01, ***p<0.001

We next tested the impact of the FAF2 knockout on neural differentiation. WT and KO cells were induced to differentiate with RA and fixed for immunofluorescence for Tubulinβ-3 (TUBB3), a neuronal marker at 3 days post induction. KO cells had significantly reduced TUBB3 fluorescent intensity compared to WT cells indicative of inhibited differentiation (**Figure 3D-F**). The blocked TUBB3 expression was also validated in whole protein lysates by western blot (**Figure 3H-I**). Additionally, neurite lengths of the few cells that did express TUBB3 in KO cells were significantly shorter when compared to WT neurons (**Figure 3G**). These experiments were repeated using a NE-4C knockdown line generated by transfecting two shRNAs and selecting with puromycin for cells that integrated the shRNA for stable expression. The shFAF2 line was kept as a polyclonal line. When the shFAF2 line was treated with RA, we also found that differentiation was blocked (**Supplementary Figure 3**).

The data above of reduced TUBB3 expression and disrupted neurite elongation indicates that knocking out FAF2 results in reduced differentiation. However, it is unclear what mechanism underlies this effect. One hypothesis is that the KO cells cannot mount a transcriptional response to promote neural differentiation when treated with RA. Bulk RNA-Seq of RA-induced cells (“RA”) that were collected 3 days post induction and untreated cells (“NO”) from WT and KO backgrounds was used to test if genes related to neural differentiation were responsive to induction. Within each genotype, differential expression analysis was used to score genes between RA and untreated conditions; the ranked genes were then used for GSEA regardless of statistical significance (**Full GSEA results in Supplementary Table 2**). Resulting terms from the WT and KO analyses were filtered by three criteria: 1) terms which were enriched in neurons relative to NPCs (normalized enrichment score (NES) > 0), 2) terms related to neural differentiation, and 3) terms that were present in both WT and KO analyses and met conditions 1 and 2. These select terms are reported in **Figure 4A**. Both WT and KO, RA-induced cells showed transcriptional signatures of differentiation with clear shifts in relative gene expression from untreated condition to RA condition (**Figure 4B**). KO cells had enrichment of genes that called terms such as “Cellular response to retinoic acid”, “Neuron fate specification”, “Neuron fate commitment”, and “Central nervous system projection neuron axonogenesis”. Several of the genes that identified these terms were significantly differentially expressed between WT RA treated cells and KO RA treated cells, however many were not (**Figure 4C**). Of the 483 genes that were used to identify the neural gene set enrichment analysis terms in WT and KO cells, 69 were differentially expressed. Taken together, this data indicates that instructive cues were initiated in KO cells in response to RA leading to a transcriptional response reflective of a neuronal fate change (**Figure 4**), however differentiation at a morphological level was blocked (**Figure 3**).

**Figure 4.**
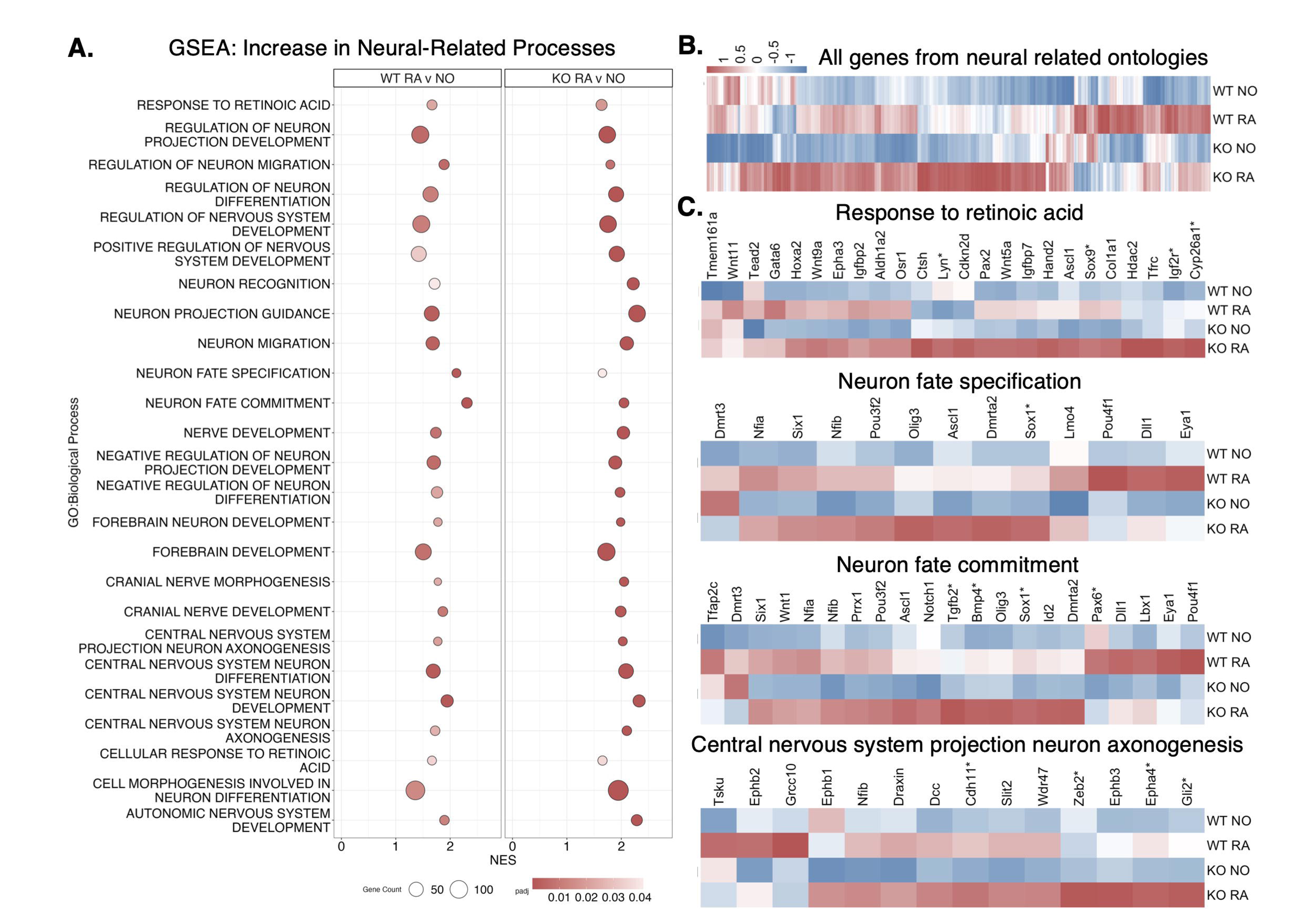
FAF2 knockout cells can mount transcriptional response to retinoic acid induced differentiation. A) Gene set enrichment analysis (GSEA) was performed between retinoic acid induced (“RA”) cells collected 3dpi and Untreated (“NO”) cells within WT or Faf2 KO genotypes. Results were filtered for positively enriched terms in both comparisons and common ontologies related to neural differentiation are shown for each genotype. B) Heatmap of all genes that identified terms in (A) C) Heatmaps of genes that identified several individual terms from (A). Asterisks (*) next to gene names indicate significant differential expression between WT neurons and KO RA-induced cells.

### Reducing ER stress is sufficient to rescue neural differentiation in Faf2 KO cells

While instructive transcriptional cues were initiated in KO cells in response to RA, neuronal differentiation was not carried out at the cellular level. This data suggests that the KO of *Faf2* and resulting excess ER stress did not inhibit the ability of cells to respond to instructive cues. We thus hypothesized that ER homeostasis may be acting as a permissive cue and that increased ER stress restricted KO cells from carrying out the morphological aspects of neural differentiation. This hypothesis was tested by reducing ER stress in KO cells and assaying if this was sufficient to rescue differentiation. To reduce ER stress, KO cells were treated with the small molecule 4-Phenylbutyric Acid (4-PBA) for 24 hours which acts as a chemical chaperone of protein folding (Kubota et al., 2006). In the 4-PBA treated KO cells relative to untreated KO cells, BIP and ATF6p50 protein expression were significantly reduced, and p-IRE expression was reduced, although this latter result was not statistically significant (**Figure 5A, B**). In contrast, p-eIF2a expression was significantly increased in 4-PBA treated KO cells relative to untreated KO cells, the importance of which is addressed in the discussion (**Figure 5A, B**). These results indicate that ER stress can be reduced in KO cells with 4-PBA.

**Figure 5.**
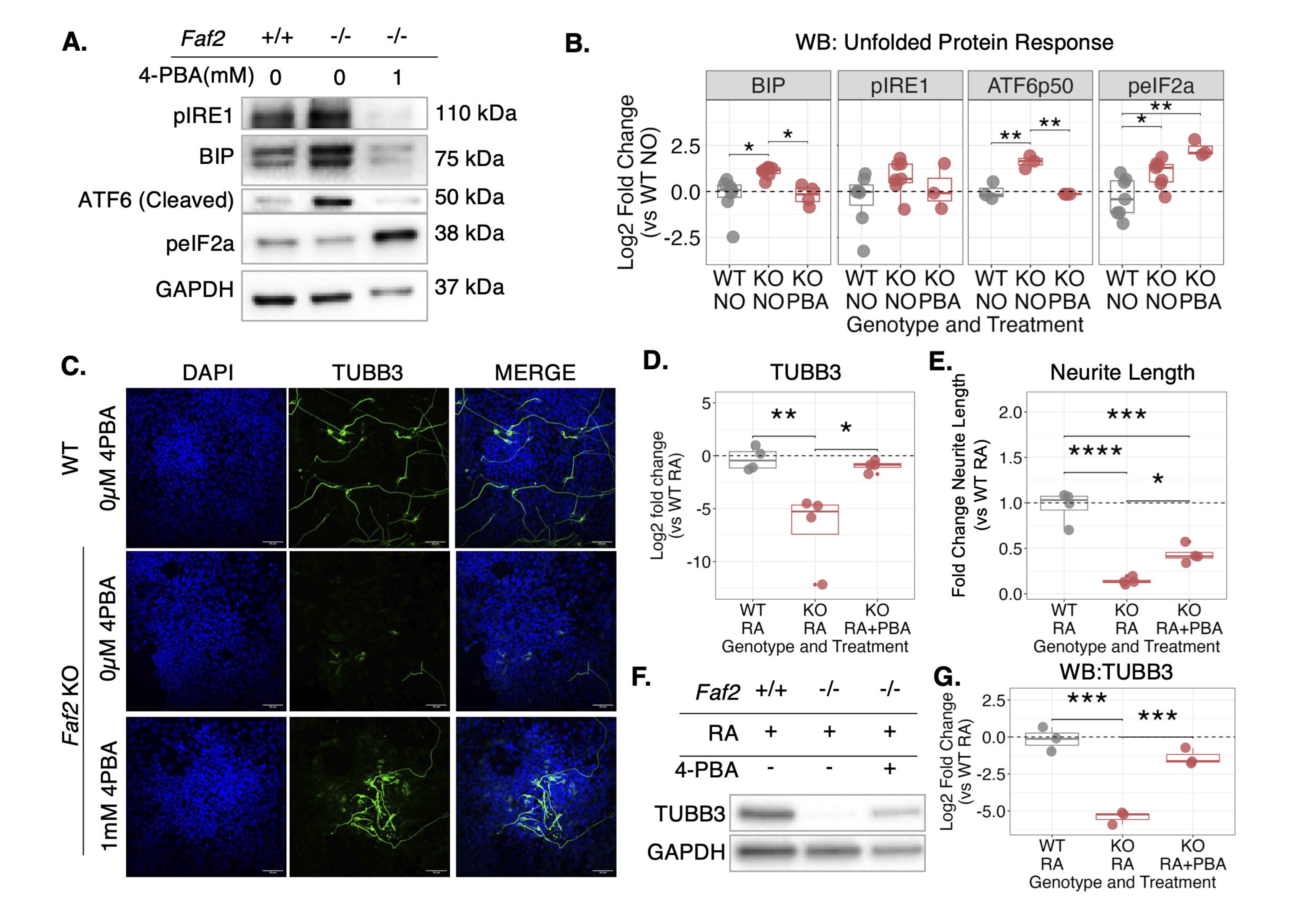
Reducing ER stress is sufficient to rescue differentiation in Faf2 KO neural progenitor cells. A) Western blot of unfolded protein response proteins (p-IRE1 110kDa, BIP 75 kDa, ATF6 50kDa, p-eIF2a 38kDa, GAPDH 37kDa) in WT, untreated KO, and 4-PBA treated KO cells. B) Quantification of western blots as in (A) C) Immunofluorescence against TUBB3 (green) in WT, untreated KO, and 4-PBA treated KO cells three days post-induction with RA. D) Quantification of TUBB3 pixel intensity. E) Quantification of neurite length. F) Western blot against TUBB3 in WT, untreated KO, and 4-PBA treated KO cells that were induced to differentiate with RA. G) Quantification of western blots as in (F). *p<0.05, **p<0.01, ***p<0.001; n= 3 biological replicates

To test if reducing ER stress is sufficient to rescue differentiation, KO cells were treated with 4-PBA for 24 hours prior to RA induction. Cells collected 3 days post induction showed significantly increased TUBB3 intensity and longer neurites in 4-PBA treated KO cells relative to untreated KO cells **(Figure 5C-G)**. 4-PBA also rescued neural differentiation in the shFAF2 knockdown cells (**Supplementary 3I-J**). This evidence indicates that in the NE-4C cell context, excess ER stress resulting from knockout of FAF2 is inhibitory to neural differentiation. Together our data implicate FAF2 as a critical player in modulating ER stress to allow neural progenitor cells to retain their competence to undergo neuronal differentiation.

## Discussion

Our study establishes that FAF2 is expressed in embryonic neural progenitor cells and is required for ER homeostasis regulation and neural differentiation. *Faf2* is relatively highly expressed when compared to genes that specify neural progenitor identity. This is not surprising as it is a ubiquitously expressed homeostasis gene necessary for regulating cell stress. In NE-4C neural progenitor cells, FAF2 protein localized to the ER as in other cell types and did not change expression, localization or intensity from cycling to differentiated states.

We find that FAF2 is a critical regulator of neural progenitor ER homeostasis. ER stress induced by thapsigargin increases FAF2 protein expression in neural progenitor cells. This indicates that FAF2 expression is responsive to ER stress and is consistent with its role in the ER associated degradation response to remove misfolded proteins (Li et al., 1993; Rutkowski and Kaufman, 2004). Furthermore, FAF2 is critical for reducing the presence of misfolded proteins as knocking out *Faf2* led to an increase in thioflavin T positive protein aggregates. FAF2 knockout cells also showed an increased signature of the unfolded protein response as the PERK and ATF6 pathways were significantly enriched at mRNA and protein levels. The loss of FAF2 function disrupts the ER associated degradation response to remove misfolded proteins and stimulates the PERK pathway which inhibits new protein production and supports autophagic removal of misfolded proteins to rebalance cellular homeostasis.

Our *Faf2* loss of function experiments revealed that FAF2 is not required for neural progenitor cell proliferation but is required for neural differentiation. *Faf2 KO* cells had greatly reduced TUBB3 expression and fewer neurons, and those cells that did differentiate had much shorter neurites. Our original hypothesis was that *Faf2 KO* cells were unable to turn on or respond to instructive cues upon retinoic acid induced differentiation. However, RNA-Seq revealed that at a transcriptional level, genes involved in neural differentiation were expressed in a similar pattern compared to WT cells. This data reveals that FAF2 is not required for interpretation of retinoic acid into downstream transcriptional programs but is needed to facilitate the conversion of those cues into the cellular behaviors reflective of neuronal differentiation such as neurite extension.

A recent study showed that excess ER stress resulting from alcohol exposure has a similar phenotype in NE-4C cells: inhibition of differentiation but no effect on proliferation (Zhang et al., 2026). Moreover, in other neural stem and progenitor cell contexts, dysregulation of ER homeostasis can disrupt neural differentiation (Kawada et al., 2014; Murao and Nishitoh, 2017). Given that FAF2 is required to regulate ER homeostasis, we hypothesized that excess ER stress could underlie the inhibition of neural differentiation in the *Faf2 KO* cells. We tested this hypothesis by treating KO cells with an inhibitor of ER stress. While BIP, ATF6 and p-IRE1 responded in a way that suggests reduced stress in *the Faf2 KO cells*, p-eIF2a was increased. Usually, treatment of cells with 4-PBA will reduce the phosphorylation of eIF2a, however in our case it does not, instead it significantly increases. Increased p-eIF2a indicates that the PERK pathway is stimulated which inhibits new protein production and supports autophagic removal of misfolded proteins to rebalance cellular homeostasis. The phosphorylation of eIF2a specifically points to attenuated protein translation (Hetz et al., 2020). Global translation of proteins and the regulation of translation are critical for the cell fate change from neural progenitor cell to neurons (Blair et al., 2017; Hoye and Silver, 2021). We hypothesize that KO cells that are already experiencing disrupted homeostasis are sensitive to the global increase in translation that is needed to support neurogenesis. Our gene set enrichment analysis of KO-RA treated cells against WT-RA treated cells shows a reduction in expression of genes in KO cells related to translation processes (Supplementary Table 2). Perhaps, p-eIF2a remains elevated to continue buffering translation and reducing misfolded protein rates, although future experiments are needed to test this idea. Regardless, 4-PBA treatment shows an overall indication of reduced ER stress and a partial rescue of neural differentiation as assessed by TUBB3 expression and neurite length. This demonstrates that the neural differentiation defect resulting from *Faf2 KO* is at least in part due to excess ER stress levels.

The partial rescue by the ER stress inhibitor suggests that FAF2 has additional functions during neural differentiation. Lipid-related processes and oxidative stress-related processes were also enriched in *Faf2 KO* cells (**Figure 2J)**. FAF2 adaptor function is critical for regulation of lipid synthesis (Ganji et al., 2023) and the maintenance of lipid droplets (Olzmann et al., 2013). While there are limited studies on the role of lipid droplet homeostasis in neural differentiation, it has been shown that reducing lipid droplet availability in adult neural stem cells can reduce neural differentiation (Ramosaj et al., 2021). Without FAF2, we hypothesize that lipid synthesis is reduced which could contribute to extra stress on neural progenitor cells and induction of transcriptional programs to increase lipid processes through the ATF6 branch of the unfolded protein response (Hetz et al., 2020). Furthermore, FAF2 can regulate lipid synthesis which is necessary for providing lipids to act as building blocks for cell membranes and to serve as signaling molecules. Studies show that treatment of neural progenitor cells with a variety of lipids can enhance neural differentiation and reducing lipid synthesis through knockout of genes such as *Fasn* can lead to a reduction in neural differentiation (Cabot et al., 2025; Gupta et al., 2025; Knobloch et al., 2013). Additionally, FAF2 is required to protect lipid droplets from being broken down by the lipase, ATGL (Olzmann et al., 2013); without FAF2, lipid droplets may be abnormally broken down, releasing triacylglycerols to the cytoplasm where they can be peroxidized and increase oxidative stress. Glutathione metabolism was an enriched term in KO cells and is a major pathway that the cell uses to combat oxidative stress (Wu et al., 2004). The case for oxidative stress in neural differentiation is complex and context specific. A neuroblastoma line (SHY-SHY cells) challenged with increased oxidative stress during neural induction showed no change in cell morphology and protein markers of differentiation but transcription was significantly altered (Khavari et al., 2020). In contrast, primary neural progenitor cells treated with hydrogen peroxide showed increased pro-neural gene expression and increased neural differentiation (Pérez Estrada et al., 2014). It is known that oxidative stress and ER stress can feedback on one another, thus reducing cellular homeostasis and stability (Cao and Kaufman, 2014). Future studies are needed to determine the relative contributions of these lipid and oxidative stress-related pathways in the cellular and molecular phenotypes of *Faf2 KO* cells. Nonetheless, our current evidence supports the model that excess ER stress downstream of loss of FAF2 function is a key factor in the block of neural differentiation.

## Conclusions

Our data show that FAF2 is expressed in neural progenitor cells with a conserved function of reducing ER stress in the cell to help maintain homeostasis. *Faf2* loss of function neural progenitor cells show a signature of increased ER stress. *Faf2 KO* does not restrict the transcriptional response to retinoic acid induced differentiation, but neuronal morphology is blocked. Reducing ER stress in *Faf2 KO* cells is sufficient to rescue neural differentiation. Taken together, we have established FAF2 as a critical player in neural progenitor cells for regulating ER homeostasis to support cell competence to act on differentiation cues.

## Acknowledgements

We acknowledge the Light Microscopy Core Facility at the University of Colorado Boulder (RRID:SCR_018993) for help and advice with microscopy. This study was supported in part by the National Institutes of Health P30CA046934 funded University of Colorado Cancer Center Genomics Core, Shared Resource (RRID: SCR_021984).

## Supplementary Data

- Supplementary Table 1: Differential expression analysis results
- Supplementary Table 2: Gene set enrichment analysis full results
- Supplementary Table 3: Gene ontology Transcription Factor full results
- Supplementary Table 4: Gene ontology Transcription Factor input genes full results

## Data Availability Statement

Raw RNA-Seq files will be made publicly available on SRA and GEO.

---

**Supplementary Figure 1.**
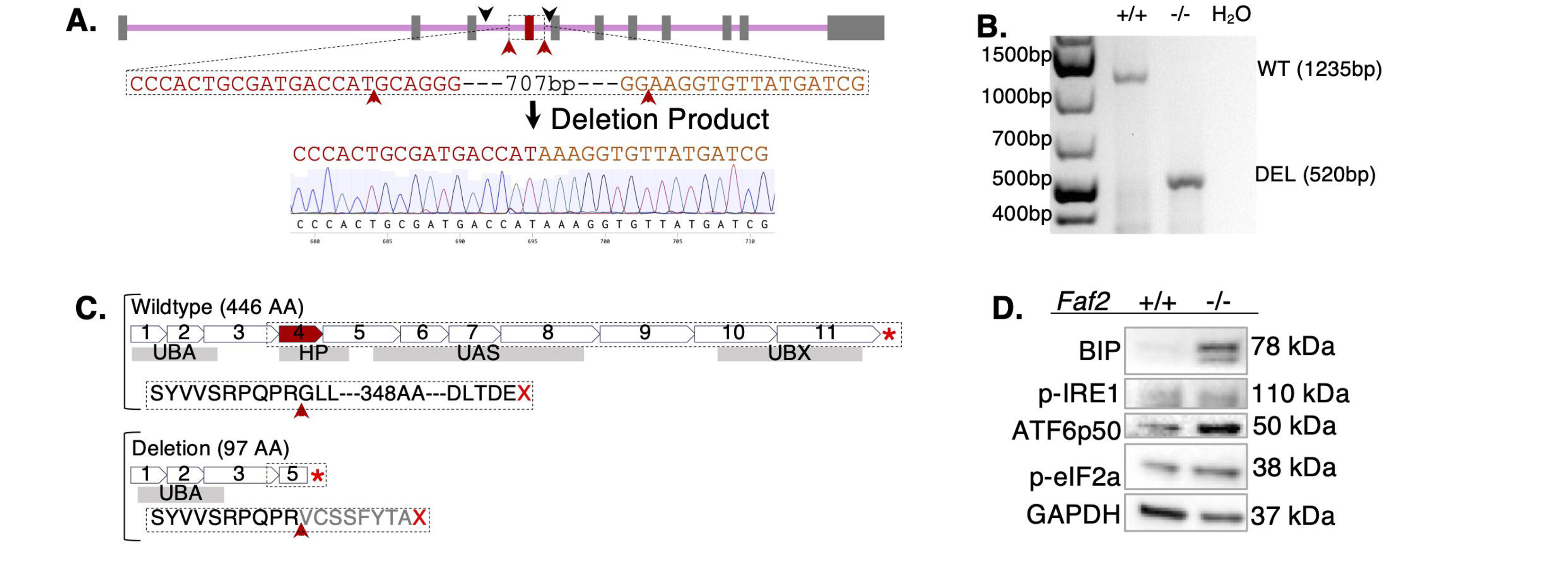
Faf2 Knockout generation. A) Gene map of *Faf2* shows exon 4 deletion target (red) between two red arrowheads. Red arrowheads indicate sgRNA target cut sites. Black arrowheads indicate primer positions for genotyping. Sequence from Sanger sequencing upstream and downstream of sgRNA sequences are shown for WT cells and in KO deletion cells. The sequencing trace is shown for the deletion product. B) 1.5% agarose gel showing genotyping results of WT and *Faf2 KO* genomic DNA. A product at 1235bp indicates WT allele and 520bp indicates deletion allele. C) Protein and domain map of wildtype FAF2 (top). Predicted protein structure for the *Faf2 KO* cells from DNA sequence in which the protein sequence is altered at the position of the red arrowhead, followed by nonsense mutation (bottom). D) Representative western blot of unfolded protein response proteins (p-IRE1 110kDa, BIP 75 kDa, ATF6 50kDa, p-eIF2a 38kDa, GAPDH 37kDa) in WT and KO cells.

**Supplementary Figure 2.**
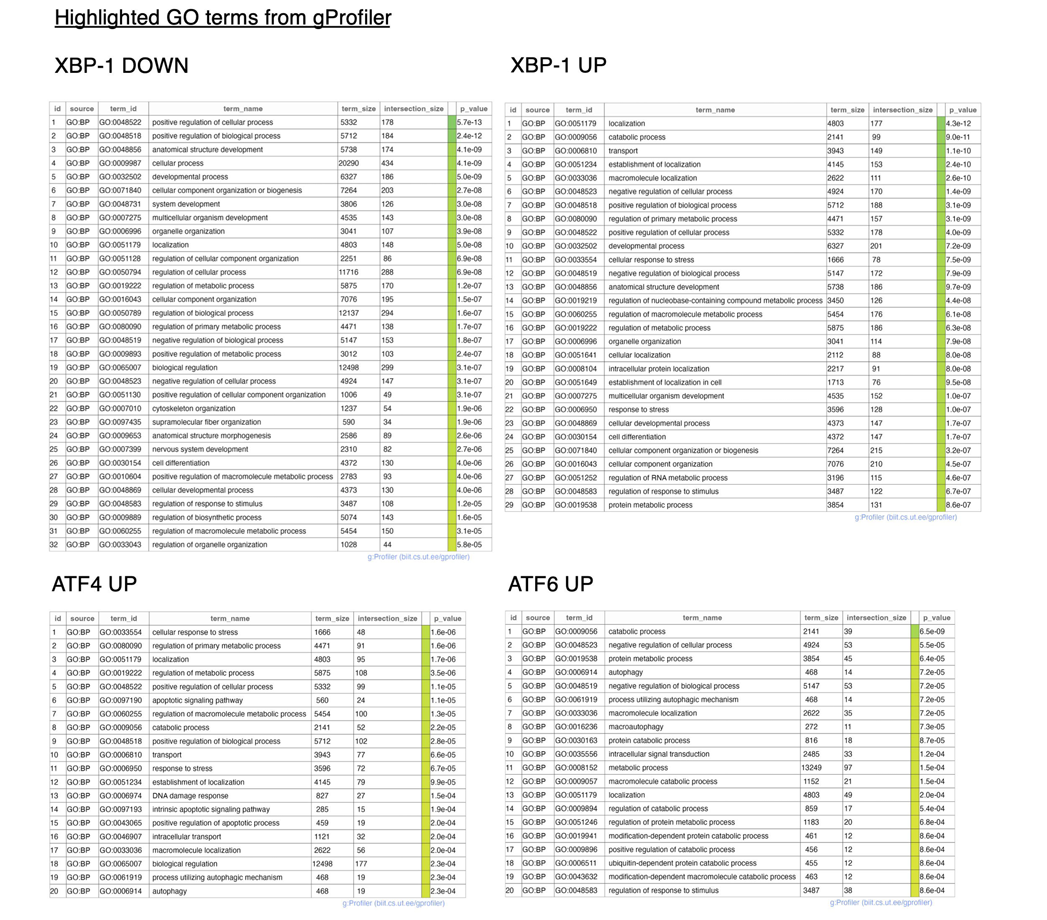
Gene ontology results from genes that call transcription factors. Genes that called transcription factors in Figure 2 were input to gProfileR for gene ontology analysis. GO: Biological Process terms are reported here for each transcription factor.

**Supplementary Figure 3.**
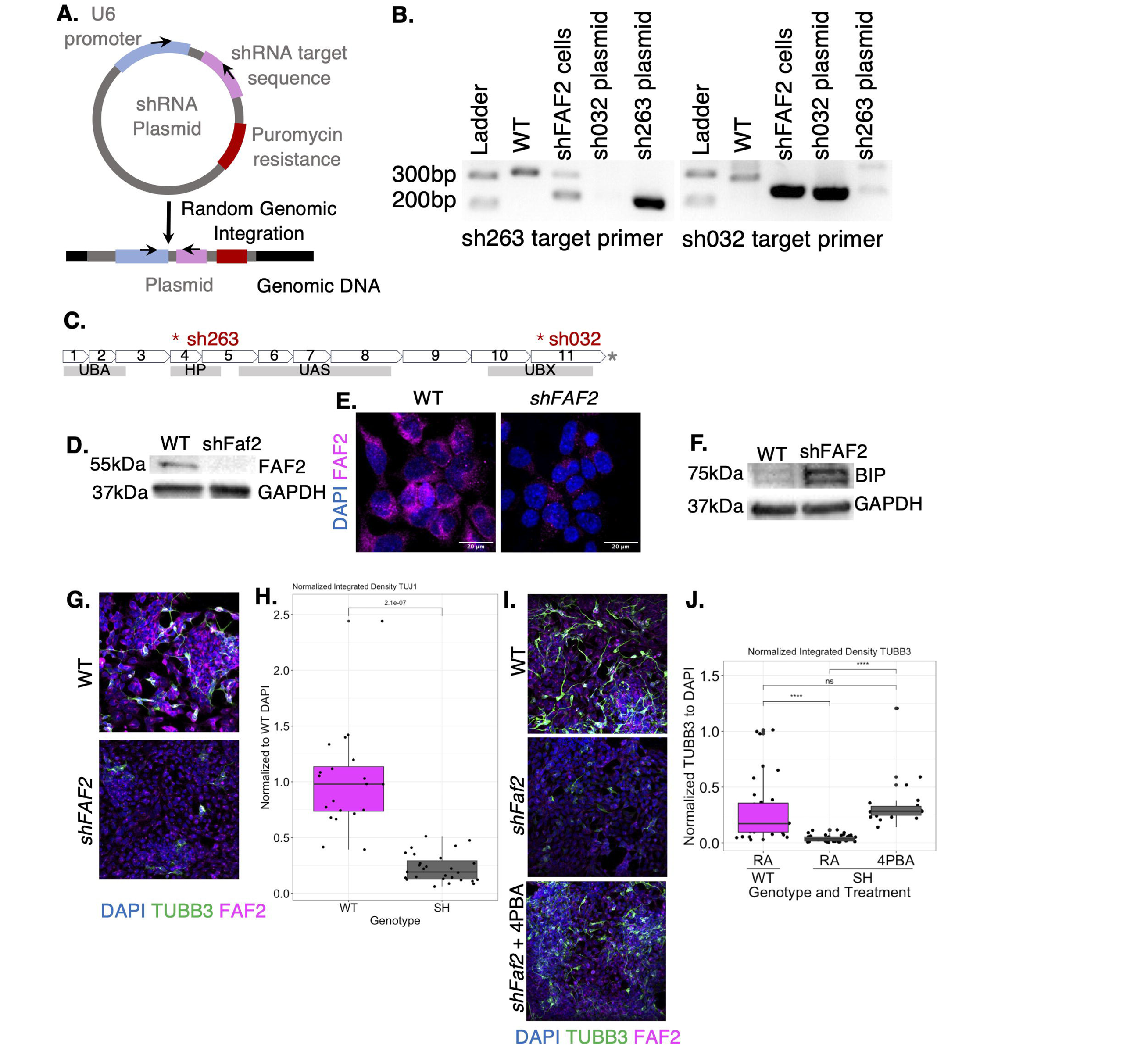
Knocking down Faf2 inhibits neural differentiation. A) Model of shRNA plasmids and strategy for detecting random genomic integration by PCR. Black arrows indicate primer locations for genotyping. B) Genotyping PCR for polyclonal shFAF2 line. Band at 230bp indicates presence of shRNA plasmid. Band at ∼300bp is an artifact band found in WT cells. Primer sets both used common forward primer on U6 promoter and shRNA target-specific reverse primers to identify the two shRNAs that were incorporated. C) Protein and domain map of where each shRNA targets FAF2. D) Western blot against FAF2 and GAPDH in WT and *shFaf2* cells. E) Immunofluorescence against FAF2 in WT and *shFaf2* cells. F) Western blot against BIP and GAPDH in WT and *shFaf2* cells. G) Immunofluorescence against FAF2 (magenta) and TUBB3 (green). Cells were induced to differentiate with RA. H) Quantification of pixel intensity from images as in (G) I) Immunofluorescence against FAF2 (magenta) and TUBB3 (green) in WT, untreated *shFaf2*, and 4-PBA treated *shFaf2* cells. Cells were induced to differentiate with RA. J) Quantification of TUBB3 pixel intensity.

